# CRISPR/Cas9-based inactivation of human papillomavirus oncogenes E6 and E7 induces senescence in cervical cancer cells

**DOI:** 10.1101/2020.04.16.044446

**Authors:** Raviteja Inturi, Per Jemth

**Affiliations:** Department of Medical Biochemistry and Microbiology, Uppsala University, BMC Box 582, SE-75123 Uppsala, Sweden

## Abstract

Human papillomaviruses (HPVs) such as HPV16 and HPV18 can cause cancers of the cervix, vagina, vulva, penis, anus and oropharynx. Continuous expression of the HPV viral oncoproteins E6 and E7 are essential for transformation and maintenance of cancer cells. Therefore, therapeutic targeting of E6 and E7 genes can potentially be used to treat HPV-related cancers. Previous CRISPR/Cas9 studies on inactivation of E6 and E7 genes confirmed cell cycle arrest and apoptosis. Here we report that CRISPR/Cas9-based knockout of E6 and E7 can also trigger cellular senescence in HPV18 immortalized HeLa cells. Specifically, HeLa cells in which E6 and E7 were inactivated exhibited characteristic senescence markers like enlarged cell and nucleus surface area, increased β-galactosidase expression, and loss of lamin B1 with detection of cytoplasmic chromatin fragments. Furthermore, the knockout of HPV18 E6 and E7 proteins resulted in upregulation of p53/p21 and pRb/p21 levels in senescent cells. These senescent cells were devoid of characteristic apoptotic markers and re-introduction of codon-modified HPV18 E6 decreased p53 levels. Taken together, our study demonstrates that cellular senescence is as an alternative outcome of HPV oncogene inactivation by the CRISPR/Cas9 methodology.

## INTRODUCTION

Human papillomaviruses (HPVs) are small, non-enveloped viruses with circular double-stranded DNA genome. HPV infections are mainly sexually transmitted and the virus exhibits tropism for cutaneous or mucosal squamous epithelium (1). The high-risk HPVs contribute to over 99% of cervical cancer cases and are also identified to cause anogenital cancers as well as head and neck carcinomas (2). More than 200 HPV types have been identified so far and among these the high-risk HPV16 and HPV18 cause 71% of cervical cancer cases worldwide. Cervical cancer is the fourth most frequent cancer in women with an estimated 570,000 new cases and 311,000 deaths in 2018 (3).

Progression of persistent HPV infections into tumor development is an unfortunate consequence for the host, considering that this is of no apparent benefit for the virus. Being a small DNA virus of approximately 8 kilobase pairs in genome size, HPV does not encode viral DNA polymerases but completely depends on the host cell replication machinery. A productive HPV life cycle requires initially proliferating epidermal or mucosal epithelial cells, where the viral DNA and early proteins are maintained at low levels in daughter cells. When cells lose the proliferation potential and differentiate, the viral DNA is amplified to very high levels along with increased expression of late genes. Uninfected epithelial cells stop proliferation and undergo terminal differentiation after a certain number of cell divisions. These cells are devoid of DNA replication and therefore cannot support virus replication. To counteract this problem, high-risk HPVs encode the two viral oncogenes E6 and E7, which prolong epithelial cell differentiation and maintain infected cells in DNA synthesis phase for the virus to use cell replication tools (4).

The high-risk HPV E7 protein interacts with retinoblastoma tumor suppressor protein (pRb) and its related proteins p107 and p130 through a conserved Leu-X-Cys-X-Glu (LXCXE) motif and disrupts the Rb/E2F transcriptional repressor complex (5,6). This results in release and activation of E2F family of transcription factors that induce unscheduled cell cycle progression. The E7 activated cell cycle progression results in stabilization and accumulation of p53 protein, which causes cell cycle arrest, senescence and apoptosis. The E6 protein from high-risk HPVs targets the p53 protein for proteasomal degradation by forming a trimeric complex with E6-associated protein (E6AP) (7,8). In this way, the E6 and E7 proteins conspire to induce and maintain cell transformation by interfering with p53 and pRb proteins that control cell cycle, senescence and apoptosis. Because the E6 and E7 proteins are crucial for the transformed cell, they can be exploited as potential therapeutic targets to control HPV-linked cancers.

The HPV or bovine papillomavirus (BPV) E2 protein binds to regulatory regions of E6/E7 viral promoters and represses transcription (9,10). As a result, expression of HPV E2 protein can efficiently block HPV E6 and E7 transcription and expression. Previous studies on inactivation of E6 and E7 by RNA interference or expression of the HPV or BPV E2 protein resulted in induction of senescence and/or apoptosis (11–14). However, application of HPV E2 or RNA interference methods has not been available for clinical treatment. CRISPR (Clustered regularly interspaced short palindromic repeats)/Cas9 is a dsDNA editing method where the Cas9 nuclease from *Streptococcus pyogenes*, guided by small RNAs, can be used to edit target DNA. Inactivation of E6 and E7 genes using CRISPR/Cas9 methodology has been reported to induce arrested cell growth and apoptosis but not senescence (15–17). However, application of CRISPR/Cas9 technology for treatment of cervical cancer needs to be addressed in more detail for a successful clinical application.

Here we investigated the CRISPR/Cas9 effect of HPV oncogene inactivation in cervical cancer cells. We found that CRISPR/Cas9 specific knockout of HPV18 E6 and E7 proteins rendered senescence like phenotype to cervical cancer cells. The senescence phenotype in cervical cancer cells was confirmed by increase in cell surface area, increased senescence associated β-galactosidase activity, loss of lamin B1 protein and detection of cytoplasmic chromatin fragments. We show that knockout of HPV18 E6 and E7 proteins resulted in activation of p53 and pRb pathways that led to cellular senescence. These senescent cells are devoid of apoptotic markers, and reintroduction of 18E6 protein into senescent cells decreased the p53 levels confirming the robustness of using CRISPR/Cas9 to knock out 18E6 and 18E7.

## MATERIALS AND METHODS

### Plasmids and constructs

The *S. pyogenes* Cas9 expression plasmid pX330-U6-Chimeric_BB-CBh-hSpCas9 (PX459) (Addgene plasmid 62988) was digested with BbsI and the gRNAs targeting E6 and E7 genes from HPV18 and HPV16 were designed using NEBuilder® HiFi DNA Assembly cloning kit (NEW ENGLAND BioLabs). Briefly, a 71 nucleotide single stranded DNA oligonucleotide containing a 21 base pair target sequence flanked by a partial U6 promoter sequence and scaffold RNA sequence was designed and purchased from Integrated DNA technologies. The ssDNA oligonucleotide and BbsI digested vector were mixed with NEBuilder HiFi DNA Assembly Master Mix and incubated for 1 h at 50°C followed by transformation into NEB® 5-alpha Competent *E. coli* (NEB#C2987).

The small guide RNAs (gRNA) were designed by utilizing Benchling online tool (Biology software 2019). Two gRNAs were designed against each target gene and two control plasmids were used in every experiment. The px459 plasmid expressing Cas9 alone was used as a common primary control in all experiments (Ctrl-1), whereas plasmid expressing Cas9 with gRNA against HPV16 E6 or E7 was used as secondary control in HeLa cell experiments (Ctrl-2).

A green fluorescent protein (GFP) expression plasmid was used as control for transfection efficiency and puromycin selection. For generating codon modified HPV18 E6, the nucleotide sequence of E6 gene was changed for modified codon usage while the original amino acid sequence of the E6 protein is kept intact. The codon modified HPV18 E6 gene was synthesized by GenScript Biotech (Netherlands) and the gene was cloned into pcDNA3.1 vector.

### Cell lines and transfection

HeLa (CCL-2) cell lines obtained from American Type Culture Collection were cultured in DMEM supplemented with 10% (v/v) heat-inactivated fetal bovine serum and 100 units/ml penicillin G and 100 μg/ml streptomycin solution (Gibco) at 37°C, 5% CO2. All experiments were performed within cell passage 7 to passage 17. The cells were transfected using Turbofect transfection reagent (Thermo Scientific) according to the manufacturer’s instructions.

### DNA extraction and analysis

Genomic DNA from Cas9 expressing cells was isolated using genomic DNA extraction buffer (10 mM Tris pH 8.0, 2 mM EDTA, 0.2% Triton X-100 and 200 μg/ml proteinase K). The extracted DNA was PCR amplified using primers targeting the starting and ending regions of HPV18 E6 and E7 genes. The gel purified DNA products were subjected to TA cloning reaction according to the manufacturer’s instructions, using TOPO TA cloning kit for sequencing (Invitrogen, 450030). The colonies were sequenced for DNA encoding E6 and E7 to confirm mutations introduced by CRISPR/Cas9. Primers targeting the starting and end regions of E6 and E7 DNA were used. The mutations were confirmed by sequence alignment to HeLa control DNA samples.

### Senescence associated-β-galactosidase assay

The senescence associated-β-gal assay was performed from puromycin selected transfected cells as previously described (18). Briefly, the cells were washed twice with ice cold PBS and fixed with fixation solution (2% formaldehyde (v/v), 0.2% glutaraldehyde (v/v)) for 5 min at room temperature. The fixation solution was removed and cells were washed again twice with PBS. Staining solution (40 mM citric acid/Na_2_HPO_4_ buffer, 5 mM K_4_[Fe(CN)_6_] 3 H_2_O, 5 mM K_3_[Fe(CN)_6_], 150 mM NaCl, 2 mM MgCl_2_ 6 H_2_O and 5-Bromo-4-chloro-3-indolyl β-D-galactoside or X-gal (Sigma, B4252, dissolved in dimethyl formamide) was used as 1 mg/ml stock solution was added to the plates and incubated overnight at 37°C. Next day, plates were washed twice with PBS, once with methanol and air-dried. The cells were imaged by phase contrast microscope (Zeiss Axio Observer Inverted Microscope).

### Western blotting

The Cas9 expressing cells with or without their respective gRNAs were positively selected by puromycin dihydrochloride (Sigma, P8833, dissolved in water), used at a final concentration of 1 μg/ml in HeLa cells for 5 to 8 days, until the gfp expressing control cells were completely dead. As a positive control for apoptosis, HeLa cells were treated with Staurosporine (Sigma, S4400, dissolved in DMSO) used at 0.4 μM and 0.8 μM final concentrations. The cells were lysed in buffer containing 25 mM Tris-HCl (pH 7.4), 150 mM NaCl, 1% NP-40, 1% sodium deoxycholate, 0.1 % SDS, supplemented with protease inhibitor (Complete Mini EDTA-free, Roche Applied Sciences). The lysates were incubated on ice for 20 min, sonicated and cleared by centrifugation at maximum speed for 10 min, 4°C (19). The supernatants were quantified using Bradford assay and subjected to electrophoresis using 8%-15% homemade gradient gels. The separated proteins were transferred to a 0.2 µm nitrocellulose membrane (Amersham Protran, GE Life Sciences) using wet transfer method for 2 h at 200 mA, 4°C. All membranes were blocked using Odyssey blocking buffer (LI-COR) at room temperature for 1 h. All primary antibodies, incubated overnight at 4°C and IRDye secondary antibodies (LI-COR), incubated at room temperature for 30 min were diluted in Odyssey blocking buffer. The membrane was washed three times in PBS with 0.05% tween 20 before and after the incubation with IRDye secondary antibodies. The membranes were imaged by Odyssey scanner (LI-COR) and quantified using Image Studio software, Version 5.2.5 (LI-COR).

Primary antibodies used in this study are anti-rabbit GAPDH (Santa Cruz; sc-47724), anti-mouse GAPDH (Ambion; Am4300), anti-mouse HPV18 E6 (Arbor Vita Corporation; AVC399), anti-mouse HPV18 E7 (Santa Cruz; sc-365035), anti-mouse Flag (Sigma; M2, F1804), anti-mouse p53 (Santa Cruz; DO-1, sc-126), anti-rabbit p21 (Cell Signaling; 2947), anti-mouse Rb (Santa Cruz; sc-102), anti-mouse phosphor-Rb (Santa Cruz; sc-271930), anti-rabbit PARP (Gene Tex; GTX100573), anti-rabbit Caspase 3 (Gene Tex; GTX110543), anti-goat Actin (Santa Cruz; sc-1616).

### Immunofluorescence and quantification

Immunofluorescence microscopy was performed as described previously (20). In brief, cells grown on coverslips were fixed in 4% paraformaldehyde for 15 min at room temperature followed by permeabilization with 0.1 % Triton X-100 for 15 min. Cells were washed three times with 2% BSA in PBS supplemented with 0.1%Tween 20 (PBST) and blocked with BSA/PBST for 1 h at room temperature. Cells were then incubated with primary antibodies: anti-rabbit Lamin B1 (Abcam; ab16048), anti-mouse α-Tubulin (Santa Cruz; sc-69969), anti-rabbit Phospho-Histone H2A.X (γ-H2AX) (Cell Signaling; 9718), diluted in BSA/PBST overnight at 4°C. The next day, Alexa Fluor-conjugated secondary antibodies (Life Technologies) were added after three washes with BSA/PBST solution. The cells were incubated with secondary antibodies in BSA/PBST for 1 h at room temperature and then washed three times with BSA/PBST. The DNA was labeled by incubation of cells with DAPI solution (Sigma, D9542, dissolved in dimethyl formamide) was used at a working concentration of 300 nM in PBS for 3 min, and washed three times with PBS. The cover slips were then mounted on slides with ProLong Diamond Antifade Mountant (Life Technologies) and imaged using Nikon eclipse 90i microscope. Quantification of signal intensity and cell measurements was done with NIS-elements AR software (Nikon).

## RESULTS

### CRSIPR/Cas9 disruption of HPV18 E6 and E7 genes results in knockout of E6 and E7 proteins

To study the effect of CRISPR/Cas9-based disruption of the genes encoding HPV18 E6 and E7, respectively, in HeLa cancer cell lines, we designed two gRNAs against each coding region. The gRNAs were constitutively expressed using a PX459 vector carrying the Cas9 gene and a puromycin resistance cassette. Throughout the study, we used the PX459 vector expressing Cas9 (no gRNA) with puromycin resistance as common control (Ctrl-1) and as a specificity control we used a PX459 vector expressing gRNAs targeting HPV16 E6 or E7 genes (E6Ctrl-2 or E7Ctrl-2) in HPV18 positive HeLa cells. Thus, Cas9 expression alone or Cas9 with gRNAs targeted against HPV16 E6 or E7 genes should not act on HPV18 E6 and E7 within HeLa cells due to the sequence difference between HPV16 and HPV18 E6 and E7 genes. HeLa cells were tested for the knockout of E6 and E7 proteins by extraction of whole cell protein lysates from puromycin selected CRISPR/Cas9 expressing cells with gRNAs against E6 and E7 regions of HPV18 and HPV16 types. As shown in figures 1A and 1C, western blot analysis using antibodies against HPV18 E6 and E7 proteins confirmed efficient knockout of E6 and E7 proteins in lysates with HPV18 specific gRNAs. The protein lysates expressing Cas9 without gRNA or with nonspecific gRNA were positive for endogenous E6 and E7 proteins confirming specificity of the CRISPR/Cas9 knockout. The intensity of endogenous E6 and E7 proteins from at least four different experiments were normalized to GAPDH levels and compared with Ctrl-1 as standard (Figs. 1B and 1D). To validate if the knockout of E6 and E7 proteins in HeLa cells was due to CRISPR/Cas9 cleavage of endogenous DNA at predicted E6 and E7 regions, we isolated genomic DNA from the cells and sequenced the regions coding for E6 and E7. We found that the predicted CRISPR/Cas9 cleavage sites within the E6 and E7 regions contained insertions or deletions that disrupted the functional E6 and E7 protein expression. The analyzed clones from HPV18 E6 gRNA-expressing cells contained three sequences with deletion mutations and one sequence with an insertion that resulted in frameshift mutation (Fig. 1E). The analyzed clones from HPV18 E7 gRNA-expressing cells contained deletion mutations and one sequence with an insertion. Taken together, we show that site-specific CRISPR/Cas9 cleavage efficiently removed endogenous E6 and E7 proteins from HPV18 HeLa cells (Fig. 1E).

**Fig. 1.**
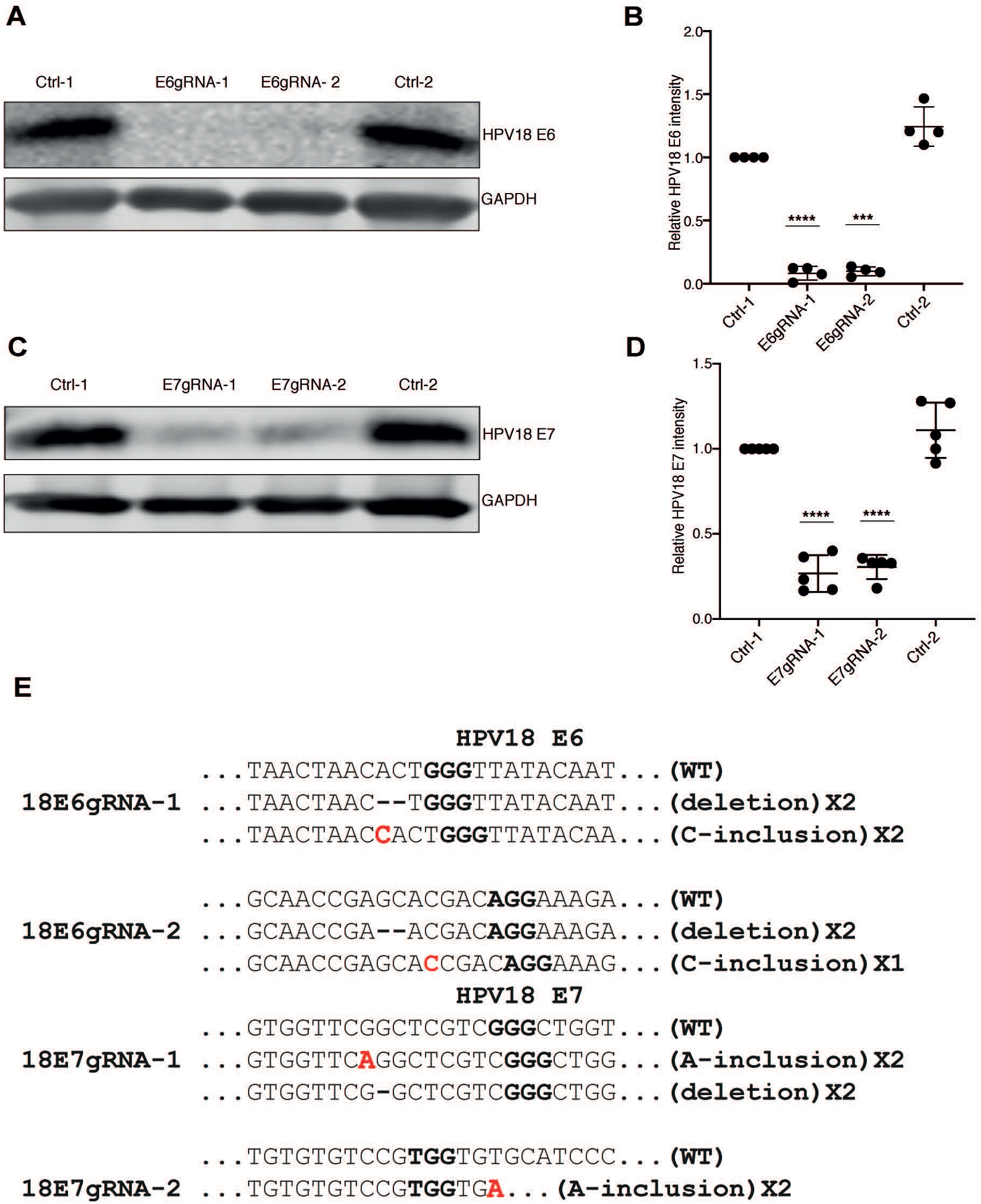
HPV18 E6 and E7 protein knockout as a result of Cas9 specific cleavage at predicted HPV genome sites. (A) Western blot analysis of puromycin selected HeLa cells expressing Cas9 (Ctrl-1), Cas9 with HPV18 E6 specific gRNAs (E6gRNA-1 and E6gRNA-2) and Cas9 with HPV16 E6 specific gRNA (Ctrl-2). The blot was probed for antibodies against HPV18 E6 and GAPDH. Representative image of four independent experiments is shown. (B) Quantification of HPV18 E6 signal intensity normalized with GAPDH and compared with Ctrl-1. (C) HeLa cells expressing Cas9 (Ctrl-1), Cas9 with HPV18 E7 specific gRNAs (E7gRNA-1 and E7gRNA-2) and Cas9 with HPV16 E7 specific gRNA (Ctrl-2) were analyzed by western blot. The blot was probed for HPV18 E7 and GAPDH antibodies. Representative image of four independent experiments is shown. (D) Quantification of HPV18 E7 signal intensity normalized with GAPDH and compared with Ctrl-1. Error bars represent mean values with standard deviation; \**P* < 0.05, \*\**P* < 0.01, \*\*\**P* < 0.001, \*\*\*\**P* < 0.0001, two-tailed student’s *t*-test was used to assess statistical significance. (E) DNA sequence alignment of HPV18 E6 and E7 sequences retrieved from HeLa cells treated with Cas9 and the respective gRNAs as indicated. The proto-spacer-adjacent motif (PAM) is denoted in bold and the red bold letters indicate the nucleotide insertions.

### HeLa cells devoid of E6 or E7 proteins exhibit senescence-associated β-galactosidase activity and altered cell morphology

Positive selections of transfected cells by antibiotic usually results in single cell colonies followed by removal of untransfected cells. To generate homogenous single cell populations with CRISPR/Cas9 expression, the HeLa cells were transfected with plasmids expressing gRNAs and Cas9 for 40 h and then selected on puromycin for over 4 days to eliminate untransfected cells. When observed under light microscope, the puromycin-selected cells with gRNAs against HPV18 E6 and E7 appeared enlarged with distinct morphology whereas the control cells with Cas9 alone or with unspecific HPV16 gRNAs formed actively growing compact single colonies (Fig. 2A). The puromycin selected cells not only failed to form single colonies, but also stopped proliferation and did not grow confluent. This was unexpected, since previous CRISPR/Cas9 studies targeting HPV E6 and E7 oncogenes have not reported this phenomenon. Senescent cells are known to exhibit distinct characteristics like increased cell morphology, elevated levels of lysosomal endogenous β-galactosidase activity detectable at pH 6.0 compared to normal growing cells (21). To further test for senescence, we performed a β-galactosidase assay on CRISPR/Cas9 treated cells. The morphologically enlarged E6 and E7 knockout cells exhibited increased blue staining, which confirmed senescence associated β-galactosidase activity. The control HeLa cells grew as compact colonies and were negatively stained for β-galactosidase activity (Fig. 2A).

**Fig. 2.**
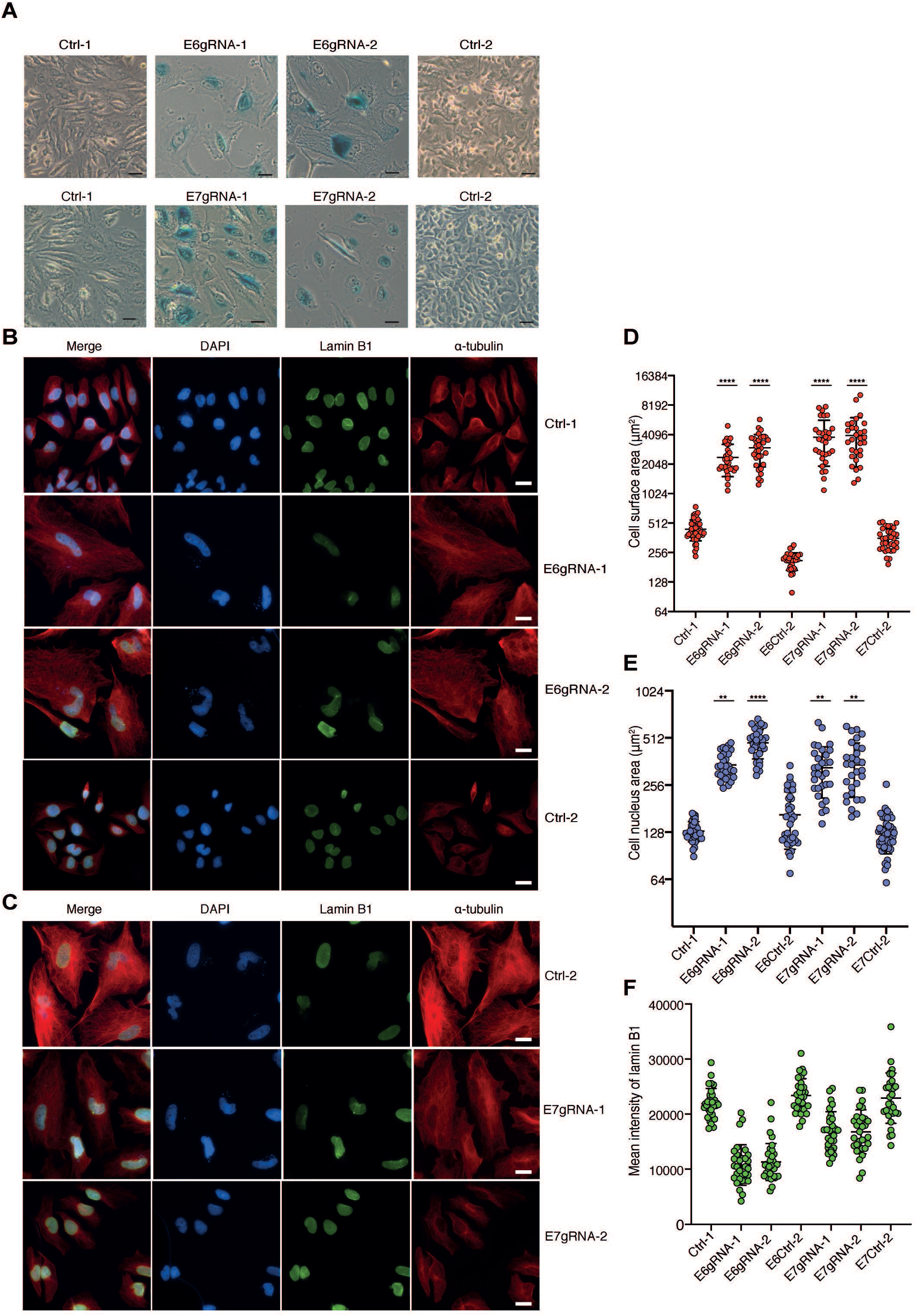
HPV18 E6 and E7 inactivated HeLa cells display senescence phenotype. (A) Light microscopic images showing senescence –associated β-galactosidase assay of HeLa cells expressing Cas9 (Ctrl-1), Cas9 with 18E6 specific gRNAs (E6gRNA-1, E6gRNA-2) or 18E7 specific gRNAs (E7gRNA-1, E7gRNA-2) and Cas9 with 16E6 or 16E7 specific gRNA (Ctrl-2) treated HeLa cells. Scale bar 100μm. Representative image of three independent experiments is shown. (B) and (C) Indirect immunofluorescence microscopy of HeLa cells expressing Cas9 (Ctrl-1), Cas9 with 18E6 specific gRNAs (E6gRNA-1, E6gRNA-2) or 18E7 specific gRNAs (E7gRNA-1, E7gRNA-2), and Cas9 with nonspecific HPV16E6 and E7 gRNAs (Ctrl-2). Shown from left to right are DAPI stained nucleus, tubulin stained cytoplasm along with lamin B1 and merge of images. Representative image of three independent experiments is shown. Scale bar 20μm. (D) The cell surface area measurements of tubulin staining cells from panel B and C plotted for the respective experiment. Mean values from three independent experiments were shown. The Y-axis is in log_2_ scale. (E) Quantitative measurements of cell nucleus surface area from DAPI stained cells from panel B. Mean values from three independent experiments were shown. The Y-axis is in log_2_ scale. (E) The lamin B1 signal from HeLa cells in panel B and C were quantified from three independent experiments and mean values are shown from each experiment. The error bars indicate mean values with standard deviation. \**P* < 0.05, \*\**P* < 0.01, \*\*\**P* < 0.001, \*\*\*\**P* < 0.0001, nested one-way analysis of variance (ANOVA) with Turkey’s multiple comparison test was used to assess statistical significance.

Enlarged cell body and irregular shape due to rearrangements in cytoskeleton is considered one of the distinct features of senescent cells. In line with that, the HeLa E6 and E7 knockout cells were detectably enlarged in size and with greater surface area than control cells. To study the extent of changes in size and morphology, we measured the cell surface area and cell nucleus surface area after performing immunofluorescence microscopy. Cells expressing Cas9 with gRNAs against E6 and E7 along with control gRNAs were permeabilized, fixed and analyzed for α-tubulin as cytoplasmic marker and DAPI staining for nucleus boundaries. By using Nikon NIS elements imaging software, individual cells were manually selected for surface area measurements. The total surface area measurements of cells as well as the nucleus were plotted separately (Figs. 2D and 2E). The total surface area of E6 and E7 inactivated HeLa cells appeared 4-fold greater when compared to control cells expressing Cas9 alone or with HPV16 E6 or E7 gRNA. Furthermore, the nucleus surface area of senescence positive cells was increased >2–fold as compared to the control HeLa cells. In summary, specific inactivation of E6 or E7 genes by CRISPR/Cas9 resulted in enhanced lysosomal β-galactosidase activity, altered cell morphology and increased total cell and nuclear surface area consistent with a senescence-like phenotype.

### HPV18 E6 or E7 inactivated HeLa cells display further hallmarks of senescence

The senescence-associated phenotype in cells is variable and heterogeneous, and there is no single universal marker to detect senescence. With increase in complexity of senescence-associated phenotypes, confirmation of senescence with more than two markers has been recommended. It is therefore crucial to use a combination of methods to confirm senescence in cells. To further validate the senescent phenotype we probed for changes in nuclear envelope protein, lamin B1 and activation of DNA damage response with detection of γ-H2AX nuclear foci, two commonly used senescence markers (22,23). Recent studies have shown that loss of lamin B1 results in nuclear changes such as appearance of cytoplasmic chromatin fragments associated with DNA damage markers like γ-H2AX (24). Hence, we tested the cells for lamin B1 levels, γ-H2AX expression and presence of cytoplasmic chromatin fragments markers by immunofluorescence microscopy.

Antibodies against lamin B1 and γ-H2AX were used to detect specific signals from CRISPR/Cas9 expressing cells and the mean signal intensity was quantified using Nikon NIS elements imaging software. The mean intensity of the quantified lamin B1 and γ-H2AX signals were plotted and shown in figures 2F and 3. As evident from the quantification, the E6 and E7 knockout HeLa cells exhibited reduced lamin B1 levels and increased γ-H2AX nuclear foci signal relative to control treated HeLa cells. Loss of lamin B1 would compromise the integrity of nucleus, which leads to detection of cytoplasmic chromatin fragments (19,20). To test this, we checked for presence of cytoplasmic chromatin fragments by DAPI stained fragments in the cytoplasm (Fig. 3). Parts of these DAPI-stained cytoplasmic fragments were found associated with γ-H2AX. On the other hand, the overall DAPI stained cytoplasmic fragments and γ-H2AX nuclear foci were relatively absent in control treated cells (Fig. 3). Taken together, these results indicate that oncogene inactivation-induced senescent cells display decrease in lamin B1 protein, increased γ-H2AX nuclear foci, and presence of DAPI stained cytoplasmic fragments associated with γ-H2AX nuclear foci.

**Fig. 3.**
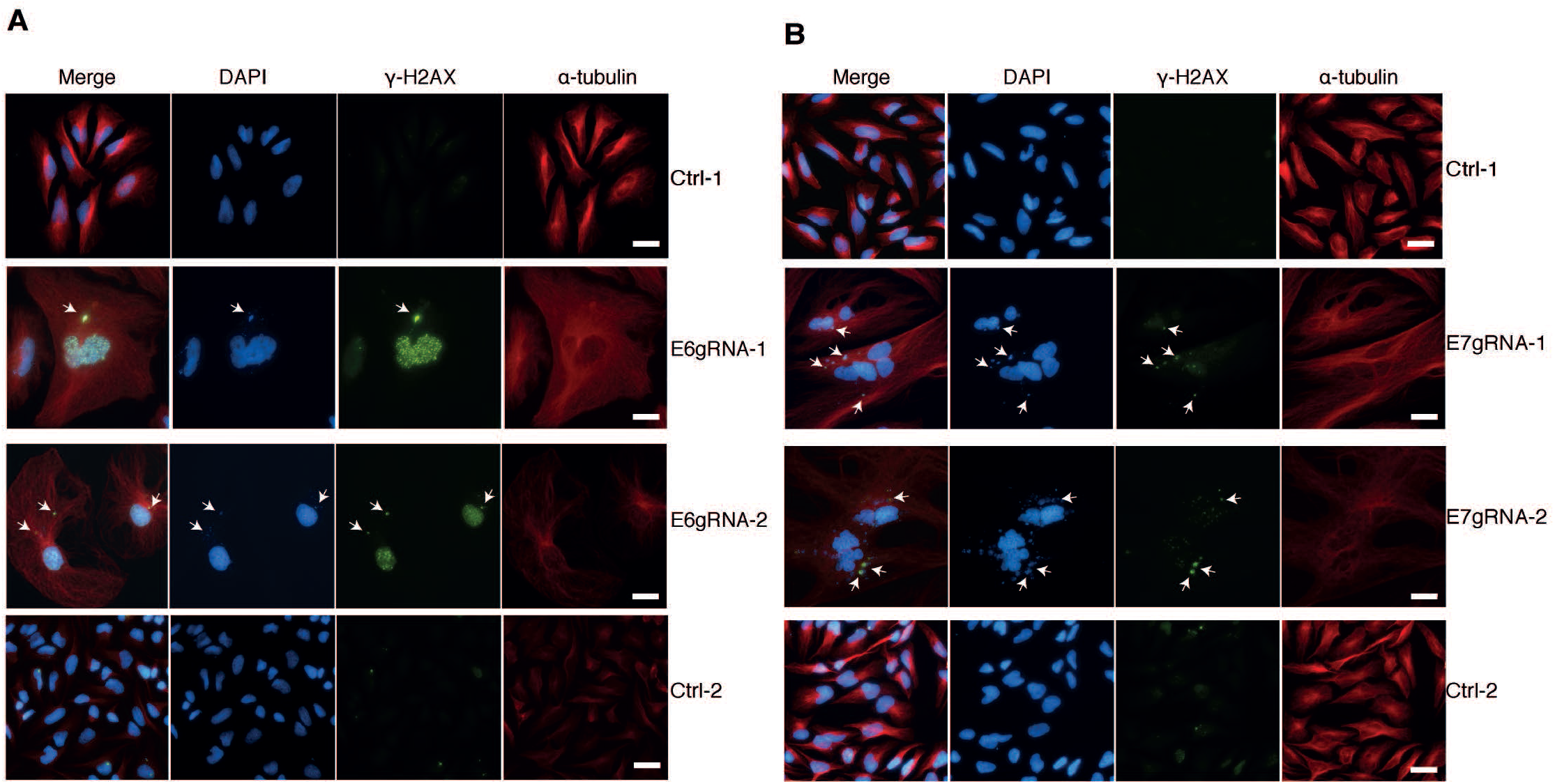
HPV18 E6 and E7 inactivated cells are positive for senescence markers. Immunofluorescence microscopy of cells with DAPI stained nucleus, γ-H2Ax and α-tubulin stained cytoplasm after transfection with Cas9 (Ctrl-1), Cas9 with HPV18 E6 specific gRNAs (E6gRNA-1, E6gRNA-2) or E7 specific gRNAs (E7gRNA-1, E7gRNA-2) and Cas9 with nonspecific gRNAs (Ctrl-2). Panel (A) represents cells with HPV18 E6 gRNAs and controls. Panel (B) shows cells with HPV18 E7 specific gRNAs and control cells. White arrows indicate cytoplasmic chromatin fragments. Scale bar 20μm. Representative image of three independent experiments is shown in both panels A and B.

### p53/p21 signaling is activated in HPV18 E6 knockout cells

The significant role of the HPV18 E6 protein is to promote proteasomal degradation of p53 protein by recruitment of the E3 ubiquitin ligase E6AP. Continuous expression of E6 protein in HeLa cells results in decreased endogenous p53 protein levels. Therefore, CRISPR/Cas9 mediated knockout of endogenous E6 protein would activate p53 signaling pathways. To test this, the whole cell protein lysates from HPV18 E6 knockout cells were made and analyzed for endogenous p53 and p21 proteins. As expected, the endogenous p53 and p21 protein levels were restored and increased expression was observed in HPV18 E6 gRNA expressed cells (Figs. 4A-4C). The Cas9 protein expression in the puromycin-selected cells was verified by using anti-Flag antibody, which detects the Flag-Cas9 protein. In the control cells the p53 and p21 protein signals remained lower confirming the presence of active endogenous HPV18 E6 protein (Figs. 4A-4C). Inactivation of HPV18 E6 restored p53 protein levels resulting in activated expression of p21 downstream signaling that could lead to cellular senescence.

**Fig. 4.**
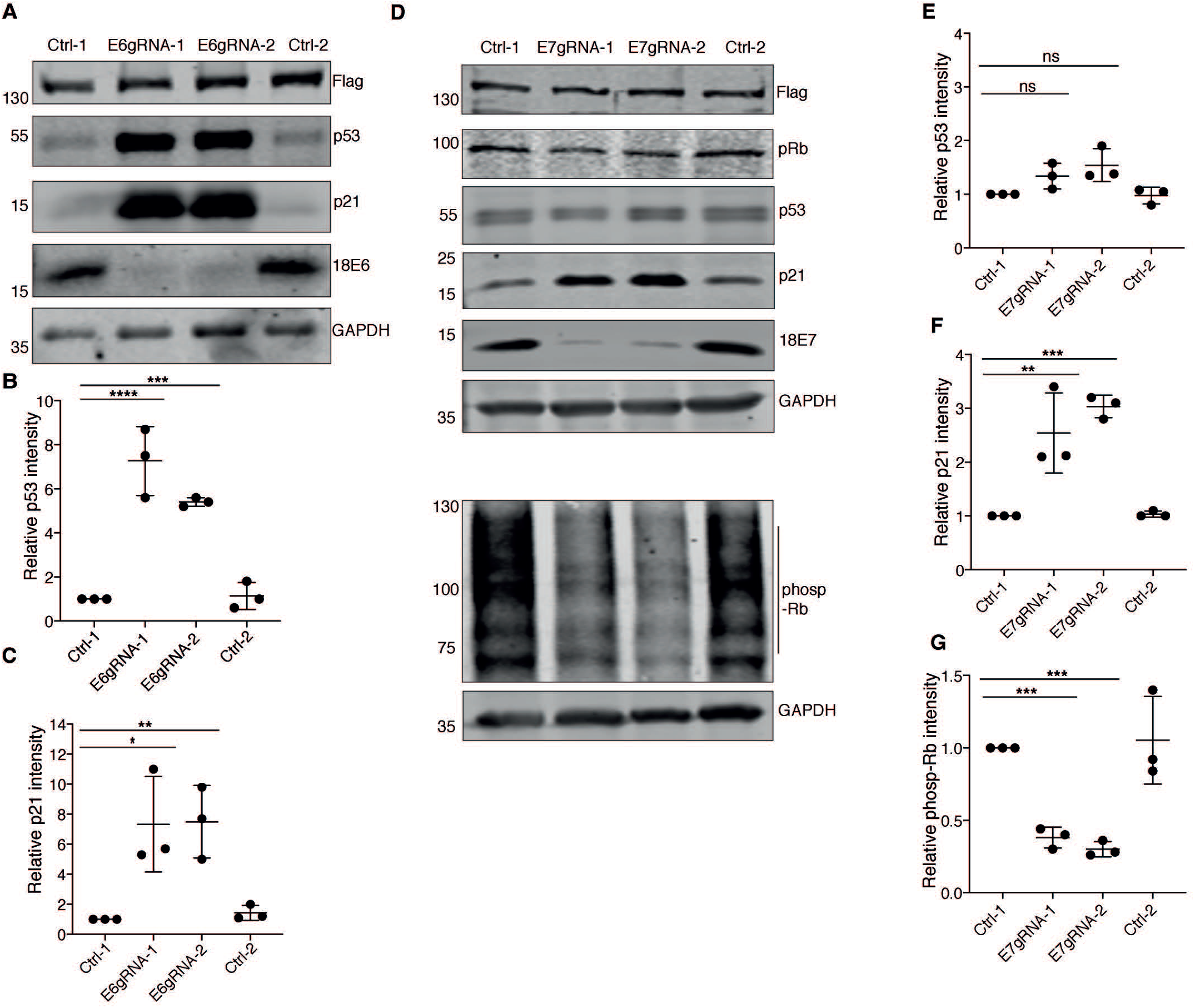
HPV18 E6 and E7 knockout restores p53 and pRb tumor suppressor pathways. (A) HeLa cells expressing only Cas9 (Ctrl-1), Cas9 with 18 E6 specific gRNAs (E6gRNA-1 and E6gRNA-2) and Cas9 with non-specific gRNA (Ctrl-2) were analyzed by western blot. The blot was probed to detect Flag-tagged Cas9 protein, p53, p21, HPV18 E6 and GAPDH. Representative image of three independent experiments is shown. Quantification of endogenous p53 and p21 levels were performed and relative intensities were plotted for the respective experiment in panel and (C). (D) Western blot analysis of protein lysates from cells treated similarly as explained in panel (A), but the gRNAs expressed were targeted for endogenous HPV18 E7 protein and the antibodies were against Flag-Cas9, retinoblastoma protein (pRb), p53, p21, and GAPDH. The lysates were re-run to detect phosphorylated-Rb (phosp-Rb). (E, F, G) The mean band intensities from three independent experiments were compared with those from Ctrl-1 and plotted versus the respective experiment. Relative intensities of (E) p53 (F) p21 (G) phosp-Rb were plotted. The error bars represent mean values with standard deviation. \**P* < 0.05, \*\**P* < 0.01, \*\*\**P* < 0.001, \*\*\*\**P* < 0.0001, two-tailed student’s *t*-test was used to assess statistical significance, ns indicate not significant.

### pRb/p21 signaling is activated in HPV18 E7 knockout cells

The HPV18 E7 protein binds to tumor suppressor protein pRb and thereby disrupts the Rb/E2F transcriptional repression. Further, the HPV16 E7 protein has been shown to destabilize and degrade Rb family proteins (25,26). In combination to disruption of Rb/E2F complex, high-risk HPV E7 proteins can directly interact and inactivate p21 and p27 proteins to maintain cell proliferation (27,28). Therefore, knockout of HPV18 E7 protein might destabilize the pRb and hence decrease the phosphorylated pRb levels in the cells. Western blot analysis of HPV18 E7 knockout HeLa cell lysates demonstrated that the pRb protein levels remained unchanged compared to control treated HeLa cells (Fig. 4D). The stable pRb protein levels in control E7gRNA-treated cells could imply pRb protein not being destabilized in HeLa cells. Notably, pRb proteins showed drastically reduced phosphorylation in HPV18 E7 knockout cells compared to control cells. The knockout of E7 protein correlates with decreased inactive pRb, which could relate to hypophosporylated Rb being available to bind E2F resulting in cell cycle arrest. The endogenous p53 protein levels in HPV18 E7 knockout cells remained relatively unchanged, but the p21 levels were elevated compared to control cells (Figs. 4D and 4E). The p53 independent increase in p21 expression could be due to the direct effect of the E7 protein on p21 protein expression. Altered pRb levels in combination with increased p21 expression can result in cellular senescence in E7 knockout HeLa cells (29). Altogether, we could confirm that knockout of either HPV18 E6 or HPV18 E7 protein levels in HeLa cells could restore endogenous p53/p21 and pRb/p21 signaling to induce senescence.

### Apoptotic markers are absent in HPV18 E6 and E7 knockout senescent cells

Previous studies on CRISPR/Cas9 application for HPV18 E6 and E7 inactivation reported apoptosis rather than senescence as downstream result. We therefore tested if the CRISPR/Cas9 inactivated HPV18 E6 or E7 cells that showed senescence (Fig. 2), also exhibited apoptosis signatures. As a positive control for apoptotic markers we treated HeLa cells with staurosporine at increasing concentrations and collected the protein lysates. The protein lysates were subjected to western blot analysis and probed for apoptotic markers using antibodies against PARP1 and caspase-3, which are cleaved to activate apoptosis (30,31). Cell lysates treated with staurosporine displayed increased levels of PARP fragments and decreased levels of full-length PARP (Figs. 5A and 5C). We did not observe any cleavage products of PARP or caspase-3 in either HPV18 E6 or HPV18 E7 knockout cell lysates which might indicate that these cells are resistant to apoptosis. In case of HPV18 E6 knockout cells we noticed a total decrease in PARP levels when compared to HPV18 E7 knockout cells. When probed for HPV18 E7 protein levels, we observed decreased endogenous E7 protein levels in lysates treated with staurosporine while the control cells retained normal E7 protein levels. The endogenous HPV18 E6 levels remained unaltered with staurosporine treatment. In conclusion, the HPV18 E6 or E7 knockout cells were negative for characteristic markers of apoptosis (Figs. 5A-5D)

**Fig. 5.**
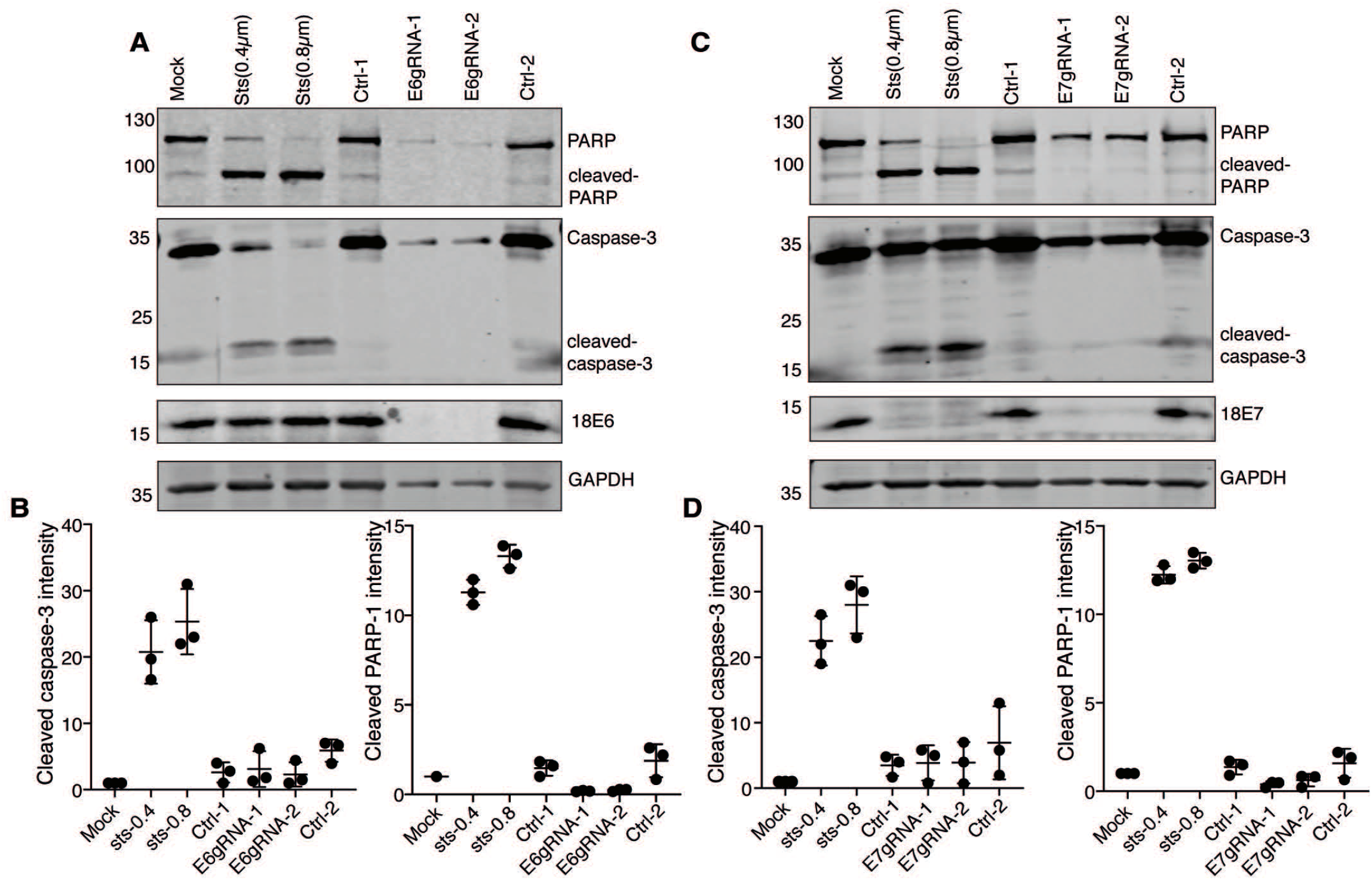
The senescent HeLa cells does not undergo apoptosis. (A) The senescent HeLa cells lysates knocked out for HPV18 E6 protein was analyzed by western blot. The blot was probed for apoptotic markers to detect PARP, cleaved-PARP, Caspase-3 and cleaved caspase-3 proteins. HeLa cells treated with staurosporine (Sts) at final concentrations of 0.4μM and 0.8μM was used as positive control for apoptosis. Representative image of three independent experiments is shown. (B) The cleaved PARP1 and cleaved caspase-3 band intensity from HPV18 E6 knockout cells were normalized with GAPDH and the mean band intensities were plotted versus the respective experiment. (C) The senescent HeLa cells lysates knocked out for HPV18 E7 protein was analyzed by western blot. The blot was probed in same way as explained in panel A. (D) The cleaved PARP1 and cleaved caspase-3 band intensity from HPV18 E7 knockout cells was normalized with GAPDH and the mean band intensities were plotted versus the respective experiment.

### Re-introduction of HPV18 E6 protein in senescent cells decreases p53 levels

To confirm the specific effect of HPV18 E6 knockout by CRISPR/Cas9, we re-introduced the HPV18 E6 protein into HPV18 E6 knockout senescent cells. Repression of endogenous E6 protein levels resulted in enhanced endogenous p53 levels (Fig. 6). At the same time, introduction of exogenous HPV18 E6 into E6 gRNA induced senescent cells lowered the total endogenous p53 levels. The alteration of endogenous p53 expression with inactivation and reintroduction of HPV18 E6 protein correlated with endogenous p21 protein expression in these cells. The switch of p53 levels with respect to E6 protein levels confirms the robustness of CRISPR/Cas9 system for specific inactivation of HPV E6.

**Fig. 6.**
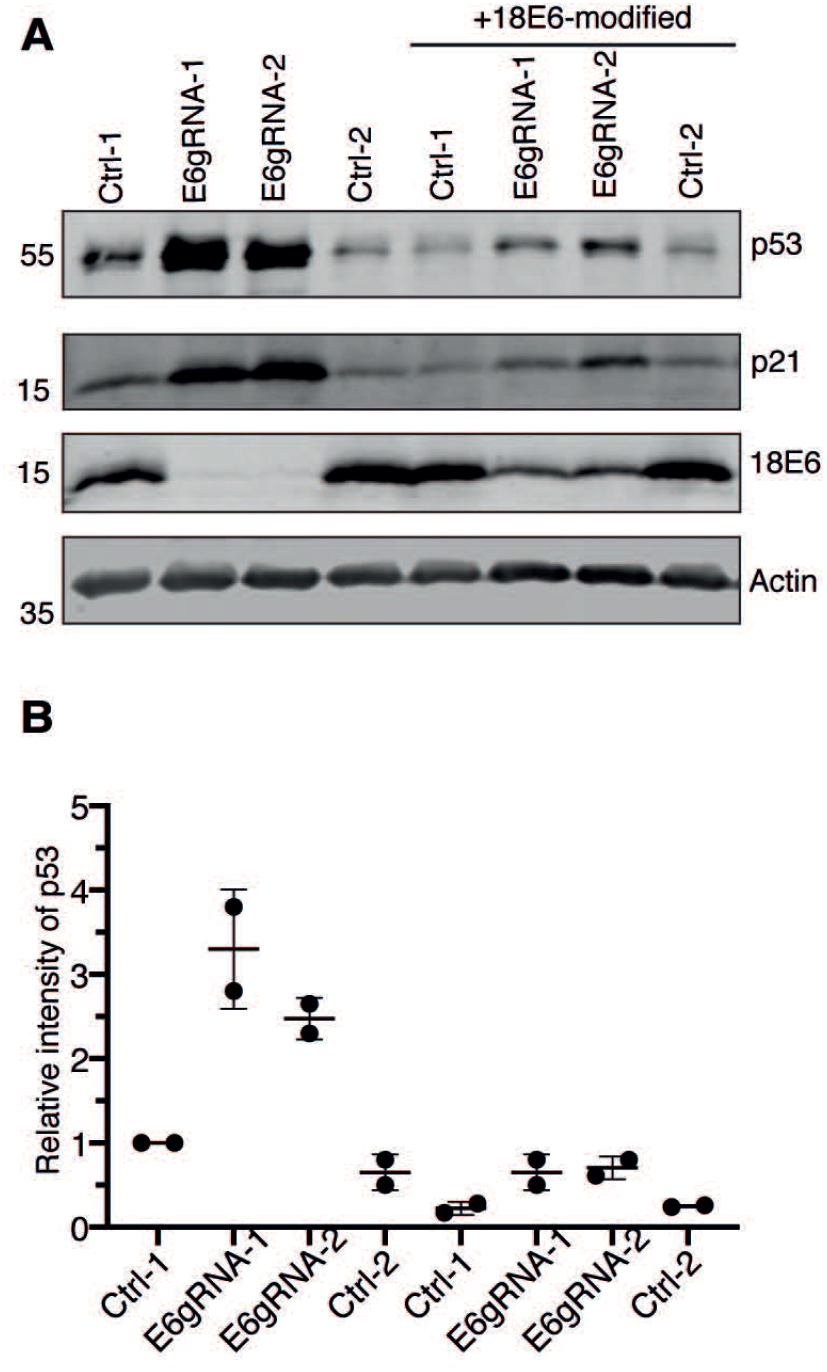
Re-introduction of HPV18 E6 in senescent HeLa cells. HPV18 E6 knock out senescent cells were transfected with either codon modified 18E6 expressing plasmid or empty plasmid followed by western blot analysis. The blot was probed for antibodies against p53, p21, 18E6 protein and actin. Representative image of two independent experiments is shown. (B) The mean p53 signal from two independent experiments was normalized with loading control actin and the relative intensity of p53 with Ctrl-1 was plotted versus the respective experiment.

## DISCUSSION

Continuous expression of high-risk HPV E6 and E7 oncogenes is essential for maintenance and survival of cervical cancer cells. The cervical cancer cells are “viral oncogene addicted” meaning that any disruption of these oncogenes is detrimental to the cells. At present, no treatment options other than surgery are available for HPV-induced cancers. Even after surgical treatment, risk of recurrence due to persistent infection compromises the chances to cure. Hence, there is an urgent need for development of treatment options to treat HPV related cancers and persistent infections. With recent advancements in DNA editing technology, CRISPR/Cas9 has evolved as a powerful genome-editing tool with precision and specificity. Using the CRISPR/Cas9 expression system we specifically knocked out integrated E6 and E7 regions from HPV18 induced HeLa cancer cells, which resulted in cellular senescence and stopped cell proliferation (32).

Integration of E6 and E7 regions within the host cell in combination with non-viral risk factors facilitate progression of HPV infections to invasive cancer. Recent work has shown that inactivation of high-risk HPV E6 and E7 genes by CRISPR/Cas9 systems restored the p53 and pRb networks resulting in induction of cell cycle arrest and apoptosis as final outcome (15–17). Studies from Jubair et al., indicated that systemic administration of Cas9/gRNAs against HPV16 E7 Caski cell and HeLa cell tumors in mice were cleared by apoptosis (17). Similarly, lentiviral transduction of Cas9/gRNA against HPV 18 E6 and E7 genes for a 10-day period induced cell death in HeLa cells (15). In line with these two studies, we transfected Cas9/gRNA expressing plasmid to target HPV18 E6 and HPV18 E7 genes and positively selected the transfected cells using puromycin. Within 4 to 6 days of selection, we noticed senescent cells with enlarged morphology instead of compact colonies as in control treated cells. Remarkably, we identified DAPI stained chromatin fragments in cytoplasm of senescent cells (Figs. 3A and 3B) and to our knowledge this the first reported example of detection of cytoplasmic chromatin fragments in HPV18 E6 and E7 knockout senescent cells. Since previous CRISPR/Cas9 studies on HPV18 oncogenes did not report senescence or assayed for senescence markers, it is difficult to exclude senescence as a potential outcome. It is possible that cells might undergo either senescence or apoptosis or both depending on the extent of intrinsic and extrinsic cellular stress. We hypothesize that the technical reasons behind the differences in alternative outcomes could be anything from differences in expression levels of Cas9, selection procedure, gRNA design, and experimental outline.

Before the advent of CRISPR/Cas9 technology, the HPV oncogene inactivation was achieved by expression of human or bovine papillomavirus E2 protein, a virus-encoded negative regulator of E6 and E7, or by RNA interference (RNAi). Inhibition of E6 and E7 expression in cervical cancer cells by E2 protein or RNAi resulted in cell cycle arrest with senescence (12–14) or apoptosis as end point (11). The induction of senescence was reported to be due to activation of dormant tumor suppressor pathways, like the p53 and pRb pathways repressed by E6 and E7 (12,33). In agreement with the above-mentioned studies, we could successfully show that CRISPR/Cas9 knockout of HPV18 E6 and E7 genes can induce cellular senescence in HeLa cells (Figs. 2 and 3). The p21 protein is one of the key mediators of therapy-induced senescence and high expression of p21 protein is often associated with senescence (29,34). With the knock down of HPV18 E6 and E7 proteins we observed the increase in p21 levels both in p53-dependent and p53-independent environments leading to senescence (Fig 4).

Cellular senescence, like apoptosis, appears to act as a defense mechanism against viruses. Expression of HPV18 E6 and E7 proteins prevents senescence in terminally differentiated cells like keratinocytes, which if untransformed undergo replicative senescence. Therefore, it is reasonable that inactivation of HPV18 E6 and HPV18 E7 genes could result in senescence. Another explanation for induction of senescence in HPV-positive cancer cells upon E6 and/or E7 repression could be due to activation of the mTOR signaling pathway. This model suggests that continued activation of mTOR signaling mediates the conversion of reversible cell growth that occurs due to repression of E6 and E7 oncogenes into irreversible growth arrest or senescence (35,36).

From a functional importance and proof of concept point, our results highlight an alternative outcome pointing senescence as permanent block of cancer cell growth. These results might be extended to develop clinically applicable viral vector or delivery systems with potential to treat HPV-linked cancers. Furthermore, senescent HeLa cells generated by inactivation of HPV18 E6 or E7 oncogenes can act as a robust cell model system to understand the details of senescence mechanisms and for screening senolytic agents to stop ageing.

## Acknowledgements

This work was funded by the Swedish Cancer Society. We thank Dr. Tanel Punga for his insightful comments on the manuscript.

